# Mechanistic insights into the Japanese Encephalitis Virus RNA dependent RNA polymerase protein inhibition by bioflavonoids from *Azadirachta indica*

**DOI:** 10.1101/2021.05.31.446388

**Authors:** Vivek Dhar Dwivedi, Ankita Singh, Sherif Aly El-Kafraway, Thamir A. Alandijany, Leena Hussein Bajrai, Mohammad Amjad Kamal, Esam Ibraheem Azhar

**Author notes:** Correspondence (E.I.A.).

## Abstract

Japanese encephalitis (JE) virus, belongs to the genus flavivirus, is a major cause of viral encephalitis, which results in neurological damage. RNA-dependent RNA polymerase (RdRp) protein is the sole enzyme responsible for viral genome replication in RNA viruses and serves as an excellent target for anti-viral therapeutic development, as its homolog is not present in humans. In this study, the crystal structure of JE RNA-dependent RNA polymerase (jRdRp) protein was obtained from protein data bank while 43 by bioflavonoids reported in *Azadirachta indica* were retrieved from the PubChem database. Following, structure based virtual screening was employed using MTiOpenScreen server and top four compounds, viz. Gedunin, Nimbolide, Ohchinin acetate, and Kulactone, with the most negative docking scores (> -10 kcal/mol) conformations were redocked using AutoDock Vina; these complexes showed mechanistic interactions with Arg^474^, Gly^605^, Asp^668^, and Trp^800^ residues in the active site of jRdRp protein against reference ligand, i.e., Guanosine-5’-Triphosphate. Furthermore, 100 ns classical molecular dynamics simulation and binding free energy calculation showed considerable stability (via hydrogen bonding and hydrophobic interactions) of docked bioflavonoids in the active pocket of jRdRp and significant contribution of van der Waals interactions for docked complex stability during simulation, respectively. Therefore, the outcome of this study predicted the substantial anti-viral activity of Gedunin, Nimbolide, Ohchinin acetate, and Kulactone against jRdRp protein and can be considered for further antiviral drug development.

## 1. Introduction

Japanese encephalitis (JE), a serious vector-borne viral infection caused by the Japanese encephalitis virus (JEV), is responsible for causing Epidemic encephalitis B — an acute infectious disease of the central nervous system, in 24 countries of Southeast Asia and the Western Pacific ^1-3^. As the transmission of JE is highly dynamic, several studies have reported substantial variation in the estimation of its global impact. For instance, a comprehensive survey estimated more than 69000 cases per year of JE in the past decade, but other estimates drastically differ incidences — from 50000 to 175000 cases per year ^4-7^. About 30 to 50 % of JE survivors have been documented with permanent neurological sequelae, imposing a heavy burden on public health and society ^6,8^. Also, a worldwide influence from JE in 2002 was estimated as 709,000 disability-adjusted life years annually^7^, advised it the one of the critical arboviral infection in humans ^2^.

The positive-sense, single-stranded RNA Japanese encephalitis virus (JEV) belongs to genus *Flavivirus*, family Flaviviridae, and has ∼ 50 nm icosahedral-shaped lipoprotein capsid, which encapsulates a 11 kb RNA genome embedded with core protein ^9^. Viral genome sequence includes 5′ and 3′ untranslated regions and a single open reading frame (ORF) with no poly(A) tail, which translates into unsegmented polyprotein of 3,432 amino acids — it further sliced and processed into three structural and seven nonstructural proteins ^10-12^. Herein, the N-terminal of the polyprotein translates for the structural proteins (capsid protein (C), membrane M protein (PrM), and envelope protein (E)), while the non-structural (NS) proteins (NS1, NS2A, NS2B, NS3, NS4A, NS4B, and NS5) are encoded by the C-terminal of the polyprotein. This maturation procedure utilizes both the viral and host proteases ^2^. Later, the seven NS proteins coordinate to form complex machinery regulating replication, cleavage of polyprotein, and fabrication of nascent viral particles ^13-15^. Among these NS protein, NS5 is the largest and the most conserved protein of the JEV which comprises an N-terminal S-adenosyl-L-methionine (SAM)-dependent methyltransferase (MTase) domain and a C-terminal RNA-dependent RNA polymerase (RdRp) region ^12,16^. The RdRp in cooperation with other NS proteins and host cell proteins participates in the viral genome synthesis by forming a membrane-bound replication complex (RC) ^17-20^. Replication process of viral genome involves the synthesis of complimentary negative RNA strand from the positive-sense single-strand RNA, which acts as a template to form double-stranded replicative form (RF). The negative strand of RF is then used as a template to produce large copies of positive strand RNA which further acts as a genomic RNA after the release from replicative intermediate (RI). The indispensable nature of RdRp in RNA synthesis for viral replication and absence of homologous protein in humans makes it a favourite target molecule for anti-viral drugs.

JEV is mainly transmitted by the mosquito *Culex tritaeniorrhynchus* in endemic regions, which prefers to breed in irrigated rice paddies ^6^, making *Culex* a primary host and reservoir to complete its sylvatic life cycle ^21^. Besides, it was predicted to maintained in an enzootic cycle in pigs and wild birds in which humans are dead-end hosts ^4^. In humans, as a neurotropic virus, it can cross blood–brain barrier (BBB) and replicate in Purkinje cells, granule cells, and pyramidal neurons of cerebrum causing a wide array of neurological complications, including acute encephalitis ^22-25^. JEV infection is more common in young children but can also infect the adults and majority of the cases shows mild or no symptoms ^26,27^. Severe clinical illness is characterized by high fever, headache, vomiting, neck stiffness, disorientation, coma, seizures, spastic paralysis, and aseptic meningitis or encephalitis ^2,28^. Although the morbidity and mortality decreased with the use of various vaccines, and several recent researchers have identified a number of compounds with anti-JEV activities, such as *N*-nonyl-deoxynojirimycin ^29^, Dehydroepiandrosterone ^30^, *N*-methylisatin-beta-thiosemicarbazone derivative (SCH 16) ^31^, Indirubin ^32^, Manidipine ^33^, Chlorpromazine ^34^, Etanercept ^35^, Minocycline ^36^, Ouabain and Digoxin ^37^, few therapies beyond intensive supportive care to treat patients with JEV are available.

To deliver the urgent need for anti-JEV therapy, traditional knowledge has long been used at the community level to limit the clinical manifestations; and hence, warrants the molecular exploration of potential novel molecules. For example, screening of natural extracts against jRdRp inhibition results in the identification of two FDA approved compounds, i.e., Ouabain and Digoxin ^37^. *Azadirachta indica* (Neem) of family Meliaceae is one such valuable traditional knowledge-based plant that exhibits medicinal properties like anticancer, anti-inflammatory, antidiabetic, antibacterial, antifungal, antimalarial, antiviral, etc. ^38^. Of note, compounds from various parts of the neem together with leaves, seeds, fruits, flowers, bark, and roots are widely used in humans because of their antimicrobial and antiviral properties ^39-42^. For instance, flavonoids from neem extracts were reported for significant inhibition of the Dengue virus type 2 ^43^. Considering the wealth of traditional knowledge and advancements in computer-aided drug discovery, we employed virtual high-throughput screening approach to screen 43 biological active bioflavonoids from neem plant against jRdRp protein. Following potential ligands were studied for binding stabilities and affinities using molecular simulations and free energy calculations, respectively to understand the detailed mechanistic mechanism and other characteristics involved in the RdRp inhibition of JEV.

## 2. Materials and Methods

### 2.1. Virtual screening and ADMET predictions

Crystal Structure of viral Japanese Encephalitis Virus RNA dependent RNA polymerase (jRdRp) protein in complex with Guanosine-5’-Triphosphate (GTP) (PDB ID: 4HDG) ^44^ solved at 2.38 Å resolution was used as the receptor molecule for structure based virtual screening with the natural compounds from neem plant (*Azadirachta indica*) at MTi-OpenScreen webserver ^45^. Herein, a total of 43 bioactive bioflavonoids reported in *Azadirachta indica* were searched in the literature and their 3D conformations were downloaded from the PubChem database (https://pubchem.ncbi.nlm.nih.gov/) ^46^. For structure based virtual screening, initially the receptor was processed by following steps: (i) removal of crystalized water molecules and heteroatoms, (ii) assignment of Gasteiger charges, (iii) addition of polar hydrogen atoms and (iv) removal of non-polar hydrogen atoms using default parameters of Dock Prep tool in UCSF Chimera-1.14 ^47^. Following, collected bioactive compounds were screened at the GTP binding pocket in jRdRp using MTi-OpenScreen webserver. After that, top four bioactive compounds with highest negative docking score were considered for further ADMET (Absorption, distribution, metabolism, excretion, and toxicity) analysis using SwissADME (http://www.swissadme.ch) ^48^ and AdmetSAR (http://lmmd.ecust.edu.cn/admetsar1/) servers ^49^.

### 2.2. Re-docking simulation and binding pose profiling

Molecular docking simulation between the jRdRp and potential natural compounds was performed by Chimera-AutoDock Vina plugin setup to decipher the most interacting residues in the active site of jRdRp against reference inhibitor, as reported earlier ^50^. Briefly, 3D structure of protein and potential bioflavonoids as ligand were subjected to minimization under the default parameters in structure minimization tool in UCSF Chimera-1.14 ^47^. Later, both receptor and ligands were primed for docking in Dock prep tool in Chimera with default parameters, where native ligand from the crystal structure was removed, polar hydrogen atoms hydrogen atoms and charges were added. Finally, molecular docking simulations were performed using AutoDock Vina ^51^ as plugin under default setting at the GTP inhibitor binding site in jRdRp protein by adjusting the grid size of 25.77 × 31.52 × 29.33 Å along both three (X, Y, and Z) axes, covering all the essential residues centre at - 38.7391, -6.2631, 29.3271 Å region, to distribute profuse space to the conformations of ligands during docking processes. Herein, at least 10 docked poses for the ligands were collected and conformation with highest negative docking score and least root mean square deviation (by default 0 in AutoDock Vina) were extracted for further binding pose analysis under default parameters using 2D interaction diagram in free academic Maestro v12.7 package (Schrödinger Release 2021-1:Maestro, Schrödinger, LLC, New York, NY, 2021). Finally, all the 3D and 2D interaction images were generated in free academic Maestro v12.7 package. Similar docking protocol was used for the GTP as reference inhibitor for comparative binding pose analysis against selected bioactive compounds.

### 2.3. Classical molecular dynamics simulation

Molecular dynamics simulation method was employed to analyse the stability and intermolecular interactions of the selected protein-ligand complexes obtained from the docking experiments. MD simulations for the docked complexes were performed under Linux environment on HP Z2 Microtower workstation using Desmond v5.6 ^52^ module of Schrödinger-Maestro v11.8 (Schrödinger Release 2018-4: Maestro, Schrödinger, LLC, New York, NY, 2018). The simulation system was prepared using the system builder module in Desmond and TIP4P model was employed for the solvation of the docked complex by placing in a 10 Å x 10 Å x 10 Å orthorhombic box. Later, the complete system was neutralized by adding Na+ or Cl− counter ions to maintain the neutrality and further minimized under default parameters using minimization tool. Finally, the whole system was subjected to simulation at a temperature of 300K and pressure of 1.01325 bar using Nose−Hoover thermostat and Martyna−Tobias−Klein method, respectively under default parameters. Each system, including reference complex, were simulated for 100 ns under default parameters with OPLS (optimized potentials for liquid simulations)-2005 force field, where a total of 5000 frames were collected at 20 ps interval during the simulation interval using molecular dynamics tool of free academic Desmond v5.6 ^52^ module in Schrödinger-Maestro v11.8 package (Schrödinger Release 2018-4:Maestro, Schrödinger, LLC, New York, NY, 2018). Finally, the generated trajectories were analysed for root-mean-square deviation (RMSD), root-mean-square fluctuation (RMSF), and protein-ligand interaction profiling with the aid of Simulation Interaction Diagram (SID) module of free Desmond v5.6 ^52^ module in Schrödinger-Maestro v11.8 package (Schrödinger Release 2018-4: Maestro, Schrödinger, LLC, New York, NY, 2018).

### 2.4. Molecular mechanics/generalized Born surface area calculation

Following MD simulation of selected protein-ligand complexes, end point based binding free energy method, i.e., Molecular Mechanics-Generalized Born Surface Area (MM/GBSA) was applied to the extracted poses at every 10 ns from each simulation trajectory to calculate the mean binding free energy under default parameters via Prime MMGBSA module of the MM/GBSA protocol in Schrödinger suite (Schrödinger Release 2018.3: Prime, Schrödinger, LLC, New York, NY, 2018). In this molecules and ions, as reported earlier ^53^. Finally, net free binding energy (ΔG) was calculated using the following equation (1).

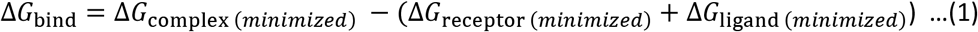

Where ΔG_bind_ denotes the binding free energy, **Δ**G_complex,_ indicates the free energy of the complex, **Δ**G_receptor_ and **Δ**G_ligand_ exhibits the energy for receptor and ligan, respectively.

## 3.1. Results and Discussion

### 3.2. Structure based virtual screening

A total of 43 bioflavonoids (Table S1) documented in *Azadirachta indica* were collected from the literature and used in virtual screening against the Guanosine-5’-triphosphate (GTP) binding pocket in jRdRp protein. All the compounds showed considerable binding affinity between -11.6 to -5.2 kcal/mol with the active residues of viral jRdRp protein (**Table S1**). Following top four compounds, i.e., Gedunin, Nimbolide, Ohchinin acetate, and Kulactone, with higher docking scores were considered for further re-docking and intermolecular interaction analysis against native crystalized ligand, i.e., Guanosine-5’-triphosphate (GTP), as reference inhibitor (**Fig. 1**).

**Figure 1.**
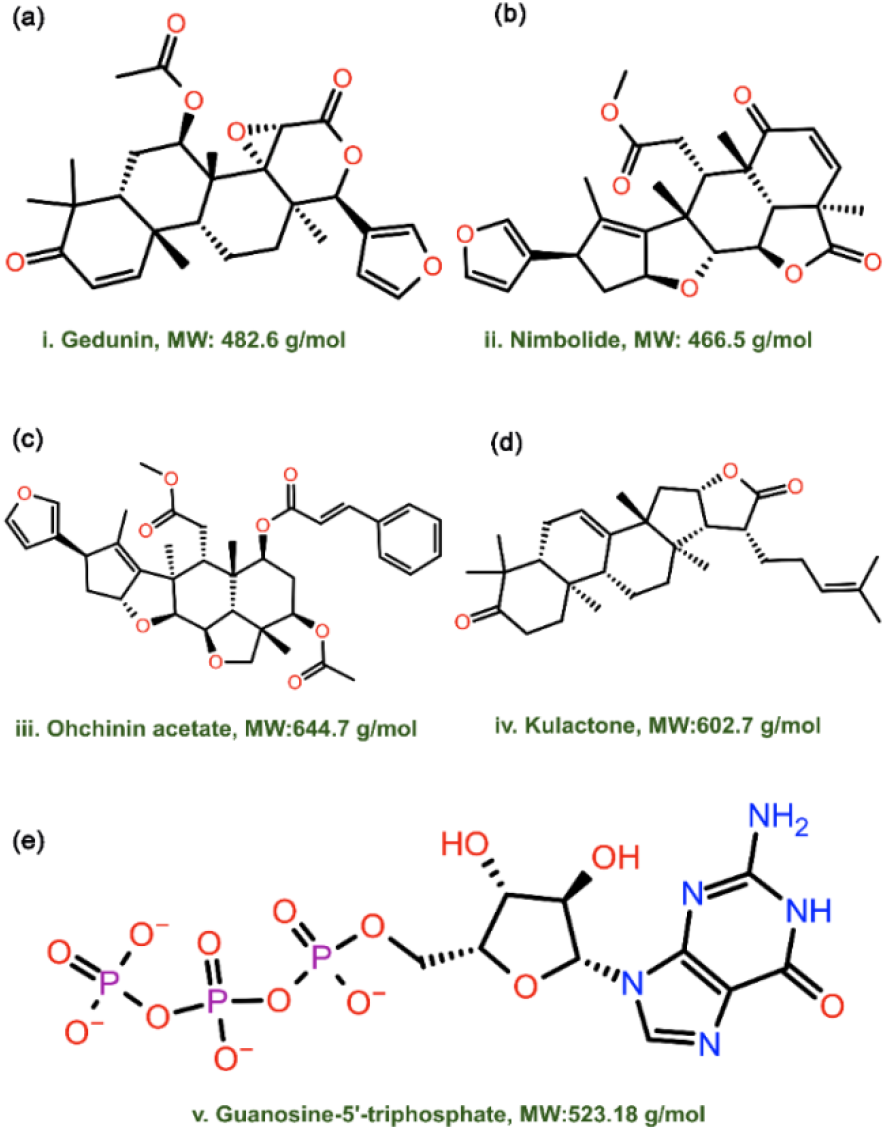
2D structural formula and molecular weight for the selected bioflavonoids, i.e., i.e., (a) Gedunin, (b) Nimbolide, (c) Ohchinin acetate, and (d) Kulactone and reference ligand, viz. (e) Guanosine-5’-triphosphate.

**Figure 2:**
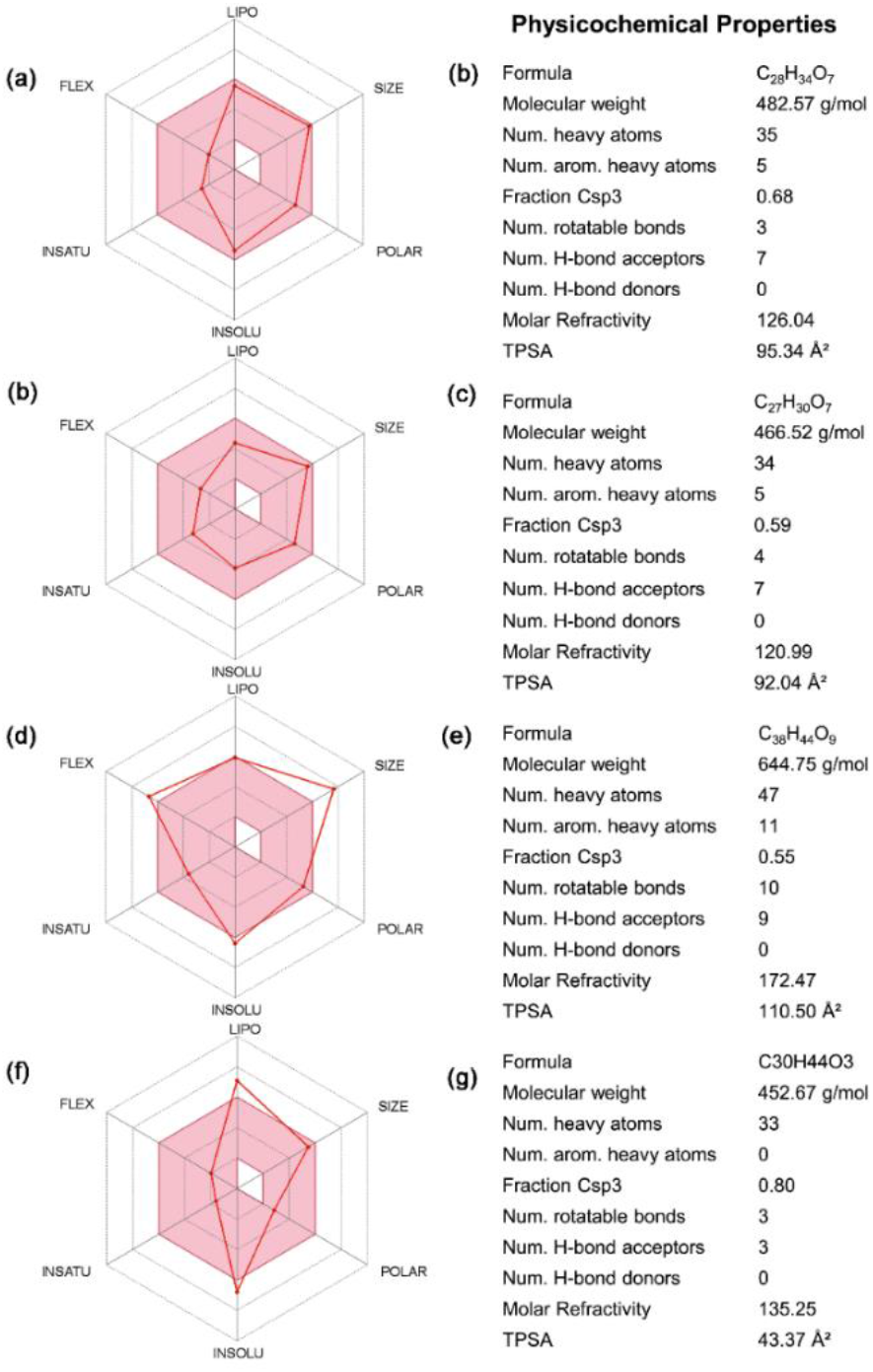
ADMET analysis for the selected bioflavonoids, i.e., (a-b) Gedunin, (b-c) Nimbolide, (d-e) Ohchinin acetate, and (f-g) Kulactone, as inhibitor against jRdRp.

### 3.3. ADME profiling

The intrinsic properties, such as drug-likeness and pharmacological characteristics, have been marked as an important factor for compounds under medical applications. Hence, ADMET properties for top four selected compounds, i.e., Gedunin, Nimbolide, Ohchinin acetate, and Kulactone, were predicted using SwissADME and admetSAR, which help to analyse the pharmacokinetic and toxic properties of compounds (**Table S2, Fig. S2**). All four selected bioflavonoids show negative AMES toxicity test and non-carcinogenic profile which reveals the non-mutagenic nature of compounds predicted using admetSAR. Cytochrome P450 2D6 (CYP2D6) is an enzyme plays vital role in the metabolism of drugs and xenobiotics and inhibition of this enzyme can lead to drug-drug interactions. Interestingly, all the selected bioflavonoids were found to be non-inhibitors of CYP2D6 using SwissADME along with other cytochromes; list of the complete ADME (absorption, distribution, metabolism, and excretion) profiling for the selected bioactive compounds is given in **Table S2**. Notably, Gedunin and Nimbolide exhibit high gastrointestinal absorption while Ohchinin acetate and Kulactone show low gastrointestinal absorption, and each of these bioflavonoids exhibit no crossing through Blood Brain Barrier (BBB). Also, Gedunin and Nimbolide shows zero violation of Lipinski’s rule of five; however, Kulactone and Ohchinin acetate shows one violation of Lipinski’s rule of five. Additionally, other four rules for drug likeness, including Ghose, Veber, Egan, and Muegge violations, showed Nimbolide as ideal candidate (**Table S2**). However, it is important to mentioned that rules for drug-likeness are not applicable to the natural bioactive compounds as these agents can be identified by the active transport system of the cells when contemplating “druggable chemical entities” ^54,55^. In addition, other properties such as pharmacokinetics and medicinal chemistry friendliness were computed for the potent compounds (**Table S2**). Conclusively, the screened compounds were suggested with ideal medical properties.

### 3.4. Re-docking and intermolecular interaction analysis

Structure based virtual screening algorithms are fast and comparatively less accurate; hence, typically best conformations of the ligands has been suggested to consider for re-docking via stringent molecular docking protocols. Thus, re-docking was performed using AutoDock Vina to collect the most suitable binding poses and conformations for the selected bioactive compounds in the active site of jRdRp protein at least docking RMSD (default 0 in AutoDock Vina) values and highest negative docking scores. Interestingly, selected bioactive compounds, i.e., Gedunin, Nimbolide, Ohchinin acetate, and Kulactone, exhibit substantial docking scores (> -10 kcal/mol) against reference inhibitor (−9.0 kcal/mol), viz. Guanosine-5’-triphosphate (**Table S3**). These results suggested the considerable binding affinity of bioactive compounds in the active pocket of jRdRp protein.

To understand the mechanistic interactions, each docked pose of bioflavonoids was studied for intermolecular interaction profile by comparison to reference inhibitor. The docked complex of jRdRp-Gedunin showed -10.4 kcal/mol docking score and formation of two hydrogen bonds (Ser^604^ and Ile^802^ residues). Additionally, hydrophobic (Leu^411^, Ala^413^, Val^414^, Ala^475^, Ile^476^, Trp^477^, Tyr^610^, Trp^800^, and Ile^802^ residues), polar (Ser^604^, Thr^609^, Asn^613^, and Ser^801^ residues), negative (Asp^541^ and Asp^668^ residues), positive (Arg^474^ and Arg^460^ residues), and Glycine (Gly^412^, and Gly^605^ residues) interactions were also noted to contribute for the stability of respective docked complex (**Fig. 2a, b**). Likewise, docking of Nimbolide with jRdRp protein exhibited -10.9 kcal/mol docking score and formation of three hydrogen bonds at Ser^604^, Ser^801^, and Ile^802^ residues. Also, jRdRp-Nimbolide complex shows additional interactions with essential residues, includes hydrophobic (Leu^411^, Ala^413^, Val^607^, Tyr^610^, Cys^714^, Trp^800^, and Ile^802^), polar (Ser^604^, Gln^606^, Thr^609^, Asn^613^, Ser^666^, Ser^801^, and His^803^), negative (Asp^668^ and Asp^669^), positive (Lys^404^, Arg^474^, and Lys^471^) and glycine (Gly^412^ and Gly^667^) interactions (**Fig. 2c, d**). Moreover, jRdRp-Ohchinin complex also presented -11 kcal/mol docking energy and noted for four hydrogen bonds (Trp^477^, Ser^604^, Ser^801^, and Ile^802^ residues), hydrophobic (Ala^410^, Leu^411^, Ala^413^, Val^414^, Ala^475^, Ile^476^, Trp^477^, Val^607^, Tyr^610^, Cys^714^, Trp^800^, and Ile^802^ residues), polar (Asn^495^, Ser^604^, Thr^609^, Asn^613^, Ser^801^, and His^803^ residues), negative (Asp^668^ and Asp^669^ residues), positive (Lys^404^, Arg^460^, Arg^474^, and Lys^47^ residues), and glycine (Gly^412^, Gly^603^, Gly^605^, and Gly^667^ residues) interactions, which were predicted to provide complex stability (**Fig.2e, f**).. Furthermore, only one hydrogen bond was observed in the jRdRp-Kulactone docked complex with -10.4 kcal/mol docking score and contribution of supplementary interactions with essential residues, such as hydrophobic (Val^414^, Phe^415^, Ala^475^, Ile^476^, Phe^478^, Tyr^610^, and Trp^800^), polar (Thr^346^, Ser^604^, Thr^609^, Asn^613^, and Ser^666^), negative (Asp^541^ and Asp^668^), positive (Arg^460^ and Arg^474^), and glycine (Gly^412^, Gly^605^, and Gly^667^) interactions (**Fig. 2g, h**). Notably, no π-π interactions and salt bridge formation were recorded in the bioactive compounds docked complexes with jRdRp protein (**Fig. 2**). Interestingly, the re-docking of reference inhibitor in the same active pocket reveals -9.0 kcal/mol and interactions with key residues, including seven hydrogen bonds formation (Lyn^463^, Arg^474^, Asp^668^, Asp^669^(2), Trp^800^, and Ile^802^), salt bridge formation (Arg^474^(3), Arg^460^ (2), and Arg^742^), hydrophobic (Met^345^, Tyr^610^, Cys^714^, Trp^800^, and Ile^802^), polar (Ser^604^, Ser^666^, Ser^715^, Ser^801^, and His^803^), negative (Asp^668^, and Asp^669^), positive (Arg^460^, Lys^471^, Arg^474^, and Arg^742^), and glycine (Gly^412^ and Gly^667^) interactions (**Fig. S1**). Of note, interesting residues were also noted in the crystal structure of jRdRp-GTP complex and essentially required to conduct the ideal reapplication of viral RNS genome ^44^. Overall, molecular contact analysis showed interaction of docked bioactive compounds with essential residues of the A, B, C, E and F motifs and the priming loop in viral RdRp protein (**Table S3, Fig. 2**). Hence, these findings suggested the screened bioactive compounds in the order of Ohchinin acetate, Nimbolide, Gedunin, and Kulactone as potential inhibitors of jRdRp by comparison to reference ligand.

**Figure 2.**
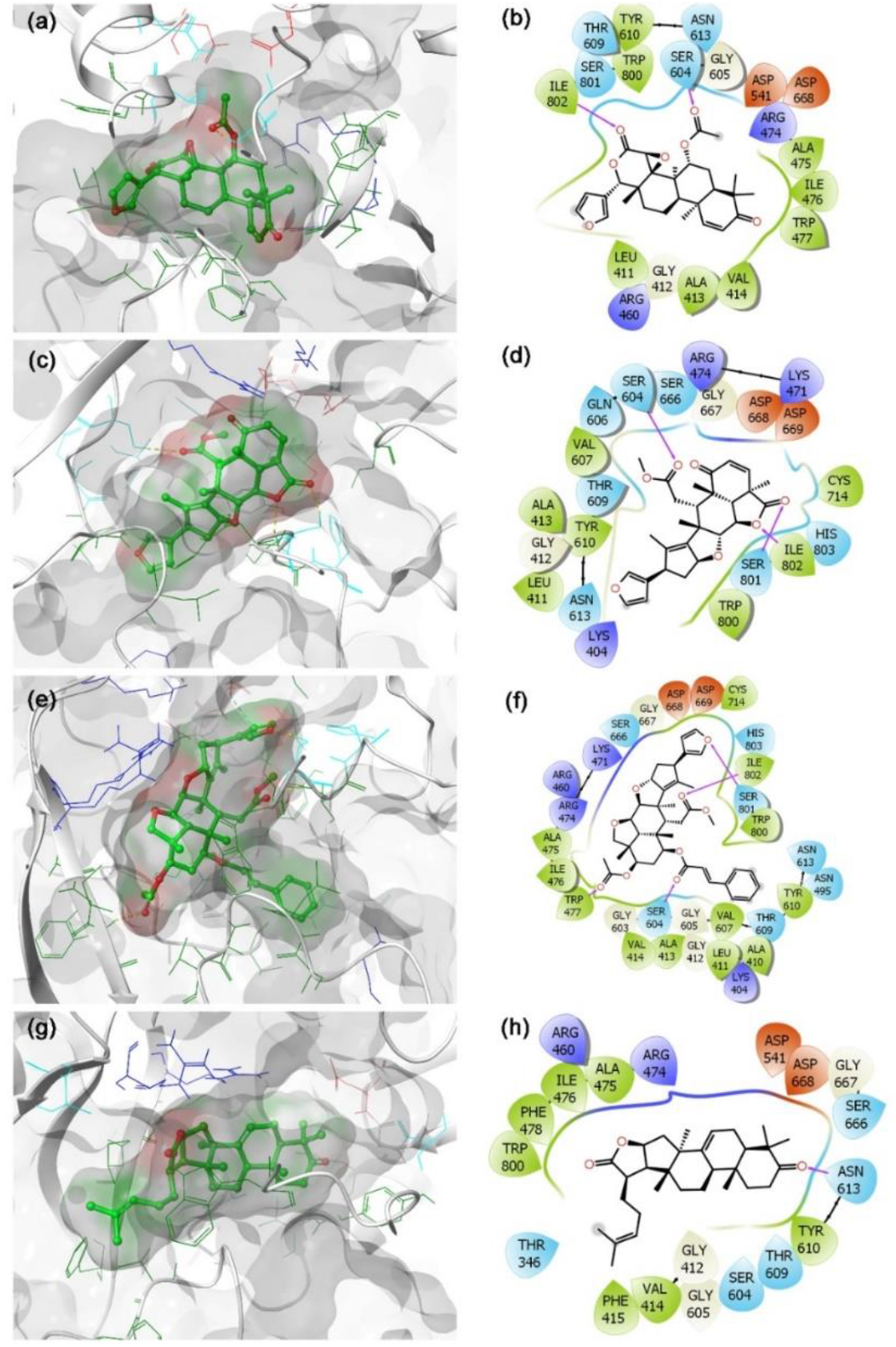
3D and 2D docked poses of the selective bioflavonoids, i.e., (a-b) Gedunin, (c-d) Nimbolide, (e-f) Ohchinin acetate, and (g-h) Kulactone, collected at 4 Å space around the ligand within in the active site of jRdRP protein. In 3D structures, protein surface and ligand surface were rendered based on the alpha-carbon and atomic charge, respectively. While in 2D maps, hydrogen bond formation (pink arrows), hydrophobic (green), polar (blue), red (negative), violet (positive), glycine (grey) interactions are logged for docked complexes of jRdRp with selected bioactive compounds.

### 3.5. Classical molecular dynamics simulation analysis

In computational drug discovery, molecular dynamics (MD) simulation is used to predict the docked complexes stability and formation of intermolecular interaction in reference to time ^56,57^. In this study, the stability of docked complexes was monitored by means of root mean square deviation (RMSD), root square means fluctuation (RMSF), and protein-ligand contact mapping, extracted from respective 100 ns simulation trajectories. Typically, RMSD and RMSF are used to observe the structural flucations which is essentially required to establish the dynamic stability of the system. Furthermore, the protein and ligands contact maps are analysed to calculate the intermolecular interaction in docked poses as function of time during simulation interval scrutinize the stability of docked ligands at the active pocket of viral protease.

#### 3.5.1. RMSD and RMSF analysis

Initially, docked complexes of potential bioactive compounds with JRdRp were analyzed for protein and ligand RMSD with respect to initial pose as reference frame (**Fig. 3**). In all the docked complexes, RMSD for jRdRp showed deviations < 2.8 Å till 60 ns, except JRdRp-Kulactone showed deviations between 4.5-5.8 Å till end of the 100 ns simulation interval. However, jRdRp docked with reference inhibitor, i.e., GTP, exhibits acceptable deviations (∼ 1.50-2.0 Å) and stability during the simulation interval (**Fig. S2**). These observations were further supported by the calculated RMSF values (<3.5 Å), except higher deviations (< 5-7.2 Å) were noted in the C-terminal of jRdRp protein docked with Nimbolide and Ohchinin acetate. Also, acceptable deviations (< 3.5 Å) were observed in the residues of jRdRp protein interacting with the respective docked ligands (**Fig. S3**). Collectively, these observations suggested the considerable rigidity and stability in jRdRp protein docked with bioactive compounds during the simulation without major structural deformations. Besides, docked bioactive compounds, i.e., Gedunin, Nimbolide, Ohchinin acetate, and Kulactone, at the active site of jRdRp protein showed considerable RMSD values till 100 ns with acceptable deviations (< 3-3.8 Å), indicates the stability of docked ligands in the active site of jRdRp protein (**Fig. 3**). Likewise, reference inhibitor, viz. GTP, also exhibits acceptable RMSD (<3 Å) fit at the active site of jRdRp protein revealed the substantial stability of respective docked complex (**Fig. S2**). Furthermore, each ligand showed acceptable RMSF values (<3 Å) fit on the protein during 100 ns simulation, inferred the significant stability of docked bioactive compounds at the active site of jRdRp protein (**Fig. S4**).

**Figure 3.**
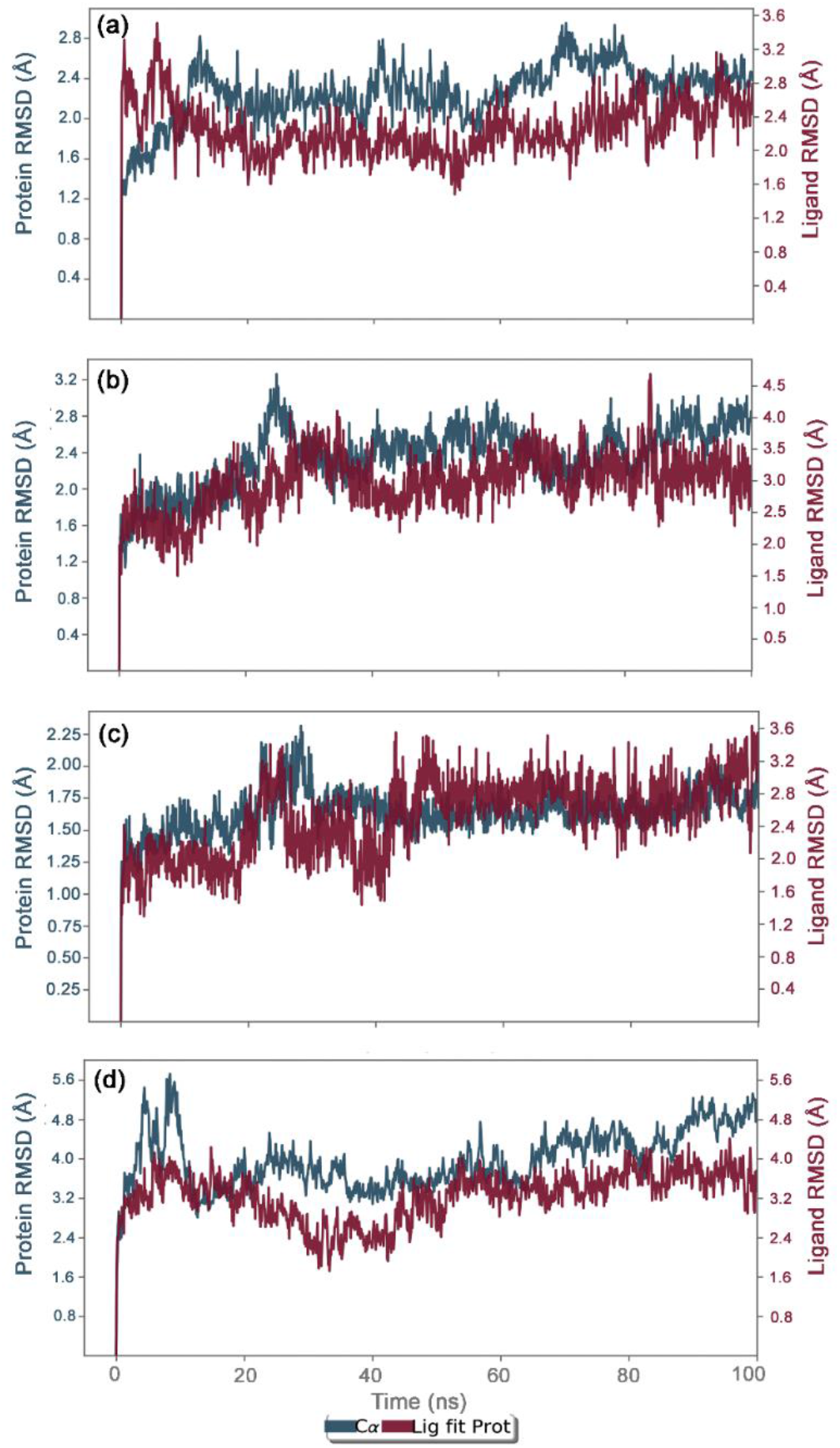
RMSD plots for the backbone atoms of jRdRp and selected bioactive compounds, i.e., (a) Gedunin (b) Nimbolide (c) Ohchinin acetate, (d) Kulactone fit on protein were extracted from 100 ns MD simulation trajectories of respective docked complexes.

#### 3.5.2. Protein-ligand interaction profiling

The docked complexes of viral protein with potential compounds were also considered for protein-ligand interaction profiling against reference complex during the course of 100 ns simulation interval in terms of hydrogen bonding, hydrophobic interactions, ionic interactions, and water bridge formation (**Fig. 4, S5**).

**Figure 4.**
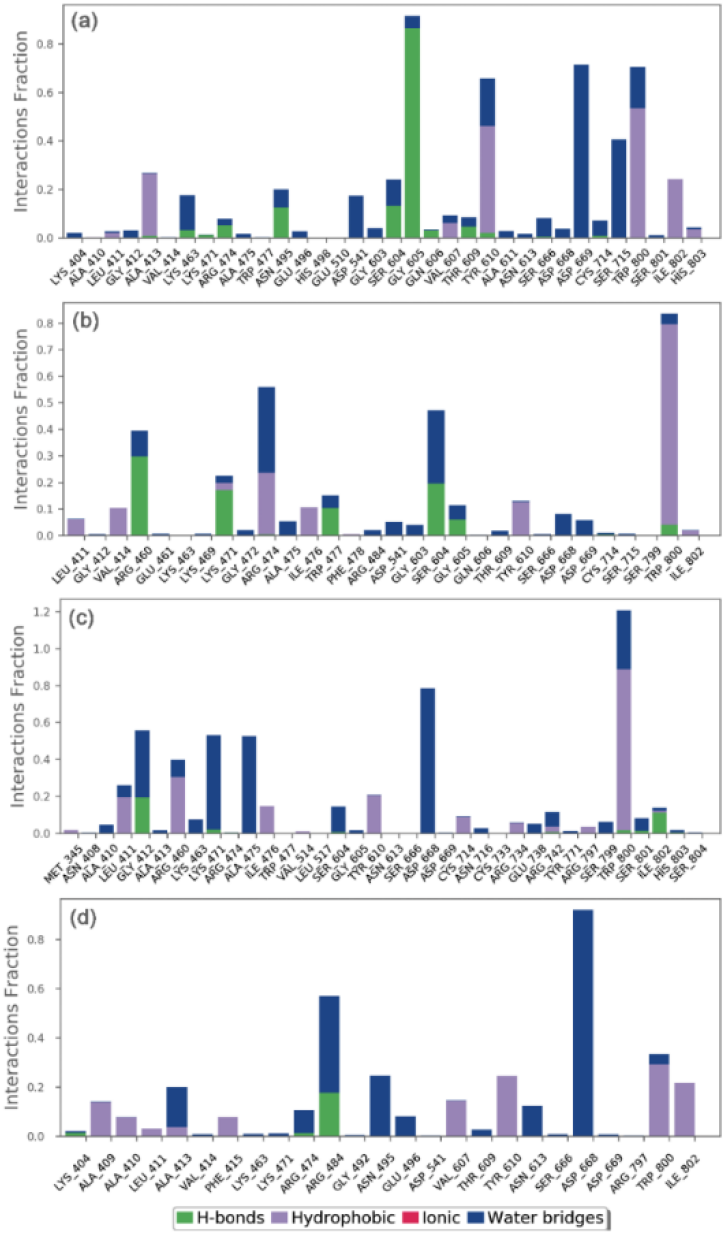
Protein-ligand interactions mapping for jRdRp docked with bioactive compounds, i.e., (a) Gedunin, (b) Nimbolide, (c) Ohchinin acetate, and (d) Kulactone, extracted from 100 ns MD simulations. Herein, values of interaction fractions >1.0 are feasible as some residues established several interactions of the similar subtype.

In the case of jRdRp-Gedunin, docked complex exhibits hydrogen bond formation for 90 % of the simulation time at Gly^605^ residue, Tyr^610^ and Trp^800^ residues noted for hydrophobic interactions for more than 50 % of the simulation time, and Asp^669^ and Ser^715^ residues participated in water bridges for more than 30 % of the MD time (**Fig. 4a**). Also, jRdRp-Nimbolide complex indicates hydrophobic interactions via Trp^800^ for 80 % of the simulation time, Arg^460^ residue was noted for hydrogen bond formation in 30 % of the simulation time, Arg^474^ and Ser^604^ residues contributed to water mediated interactions for 30 % of the total simulation time (**Fig. 4b**). Moreover, jRdRp-Ohchinin complex exhibits the most stable hydrogen bond interaction at Arg^474^ with more than 100 % of the simulation time and water bridges formation at Asp^668^ residue for 75 % of the simulation interval (**Fig. 4c**). Furthermore, significant water bridge interactions via Arg^484^ (40 %) and Asp^668^ (90 %) were noted for the jRdRp-Kulactone docked complex during 100 ns simulation interval (**Fig. 4d**). However, reference complex, viz. jRdRp-GTP, shows the most significant interaction at Arg^474^ via both hydrogen and water bridge formation for 50 % of the simulation interval (**Fig. S5**). Interestingly, the interactive residues were also observed in the respective docked complexes and essentially required for the replication of virus (**Table S2**). Hence, these results support the considerable stability of selected compounds at the active pocket of jRdRp by formation of strong hydrogen bonding and hydrophobic interactions; hence, can be used as potent inhibitors of jRdRp protein.

Additionally, intermolecular interactions between the residues of jRdRp protein and potential bioactive compounds, i.e., Gedunin, Nimbolide, Ohchinin acetate, Kulactone, and reference ligand, viz. GTP, were calculated at total 30% interval of 100ns simulation which revealed considerable binding of respective ligands with active residues (**Fig. 5, S6**). Interestingly, all the selected ligands were observed for hydrogen boning and water bridge interactions, suggested the stability of selected compounds at the active site of jRdRp protein. Hence, based on 100 ns molecular dynamics simulation analysis, docked complexes can be arranged in order of stability, i.e., jRdRp-Ohchinin acetate, jRdRp-Gedunin, jRdRp-Kulactone, and jRdRp-Nimbolide.

**Figure 5.**
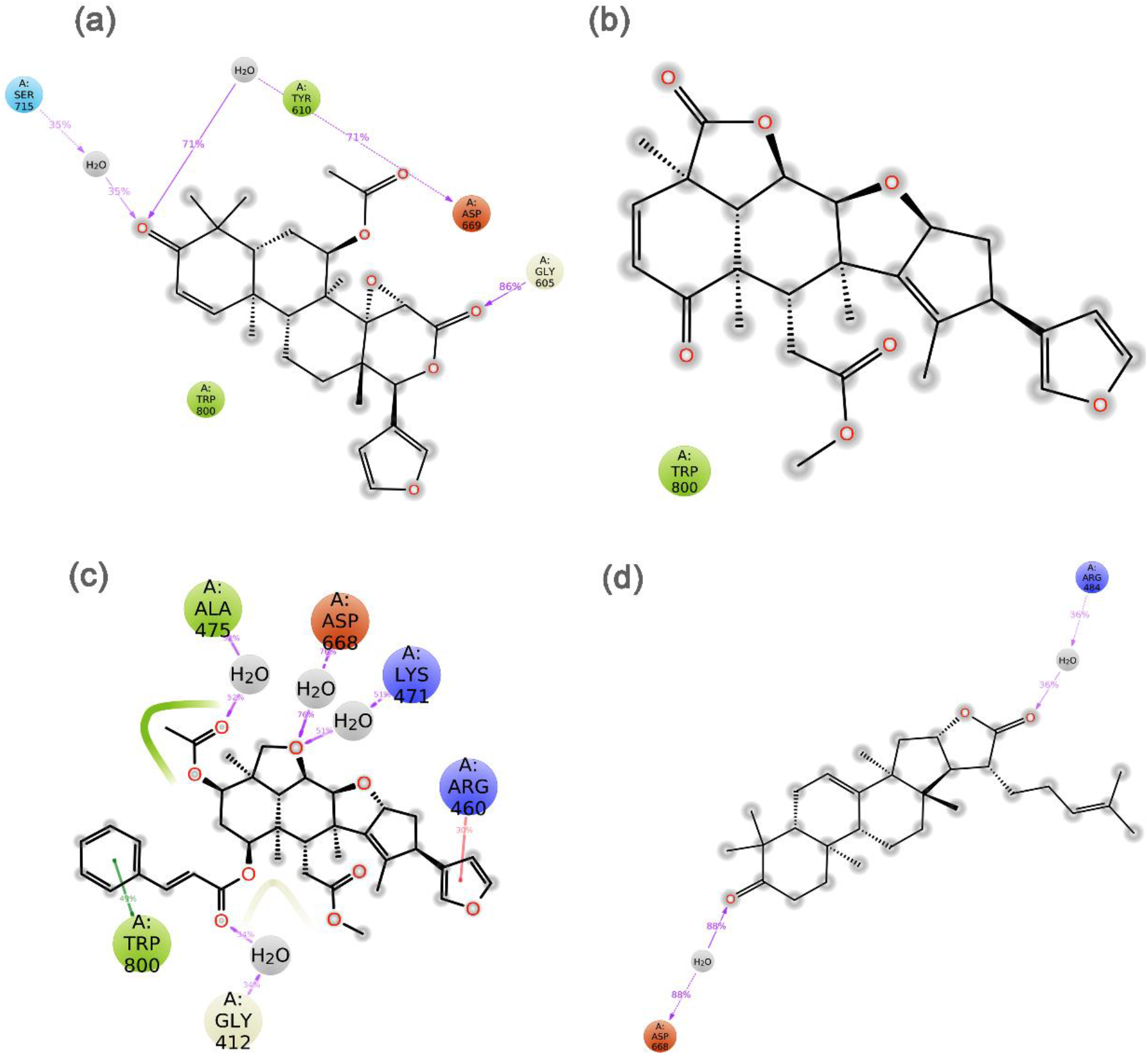
Schematic representation for interaction profile of jRdRp protein docked with bioactive compounds, i.e. ((a) Gedunin, (b) Nimbolide, (c) Ohchinin acetate, and (d) Kulactone, extracted at 30 % of total 100 ns simulation interval.

### 3.6. Binding free energy analysis

The binding affinities of the four selected protein-ligand complexes were estimated using MM/GBSA approach. Herein, poses of jRdRp-bioactive compounds, i.e., Gedunin, Nimbolide, Ohchinin acetate, and Kulactone, were extracted from 100 ns MD simulation trajectory at every 10 ns interval for average free binding energy calculations against reference complex, viz. jRdRP-GTP complex (**Table S4, Fig. 6, S7**). All the docked complexes of bioactive compounds with jRdRp exhibits significant binding free energy (> -50 kcal/mol). Of note, highest ΔG_Bind_ (−61.13 ± 3.26 kcal/mol) was recorded for jRdRp-Nimbolide complex against jRdRp-GTP (ΔG_Bind_ = -78.9±9.05 kcal/mol). Moreover, calculation of dissociation energy components for each complex exhibits substantial support of ΔG_Bind Lipo_, and ΔG_Bind vdW_ in the stability of docked complexes while ΔG_Bind Solv GB_ was observed in contribution for instability of the respective complexes (**Fig. 6, S7**). Conclusively, results suggested the considerable stability of docked bioactive compounds in via formation of van der Waals interactions with the essential residues in the active site of jRdRp protein.

## 4. Conclusion

The role of RNA dependent RNA polymerase (jRdRp) in the replication and survival of RNA genome viruses inside a host and absence of human homolog makes it an attractive molecular target for anti-viral drugs. Besides, board range of secondary metabolites as natural bioactive compounds in *Azadirachta indica* have been reported with medicinal benefits. Hence, this study evaluated the potential of known bioactive compounds in *Azadirachta indica* as potential inhibitors of jRdRp protein using complex molecular simulation, drug likeness profiling, and end-point binding free energy calculations. Herein, among the screened 43 bioactive compounds at the active site of jRdRp protein, four bioflavonoid compounds, viz. Gedunin, Nimbolide, Ohchinin acetate, and Kulactone, with substantial docking score and druglikeness were considered for further mechanistic inhibition analysis of jRdRp protein. The analysis of docked complexes showed that selected bioactive compounds occupied the active site by both hydrogen bond, hydrophobic interactions, and other considerable intermolecular interactions with essential residues of jRdRp protein. Moreover, molecular dynamics simulation and free energy calculation further revealed the substantial stability and contribution of van der Waals interaction in the stability of respective docked complexes, respectively. In conclusion, these results supported the Gedunin, Nimbolide, Ohchinin acetate, and Kulactone as acceptable inhibitors of jRdRp protein and advised to study for further in vitro and in vivo experiments to develop potential treatment against Japanese Encephalitis Virus.

## Conflict of Interest

The authors declare that there is no conflict of interest.

## Acknowledgement

Author, Ankita singh acknowledge the Director, AIRF, Jawaharlal Nehru University, New Delhi, India for giving access to Schrodinger program.

## Supplementary Section

Table S1-S4

Figure S1-S7

---

**Figure S7.**
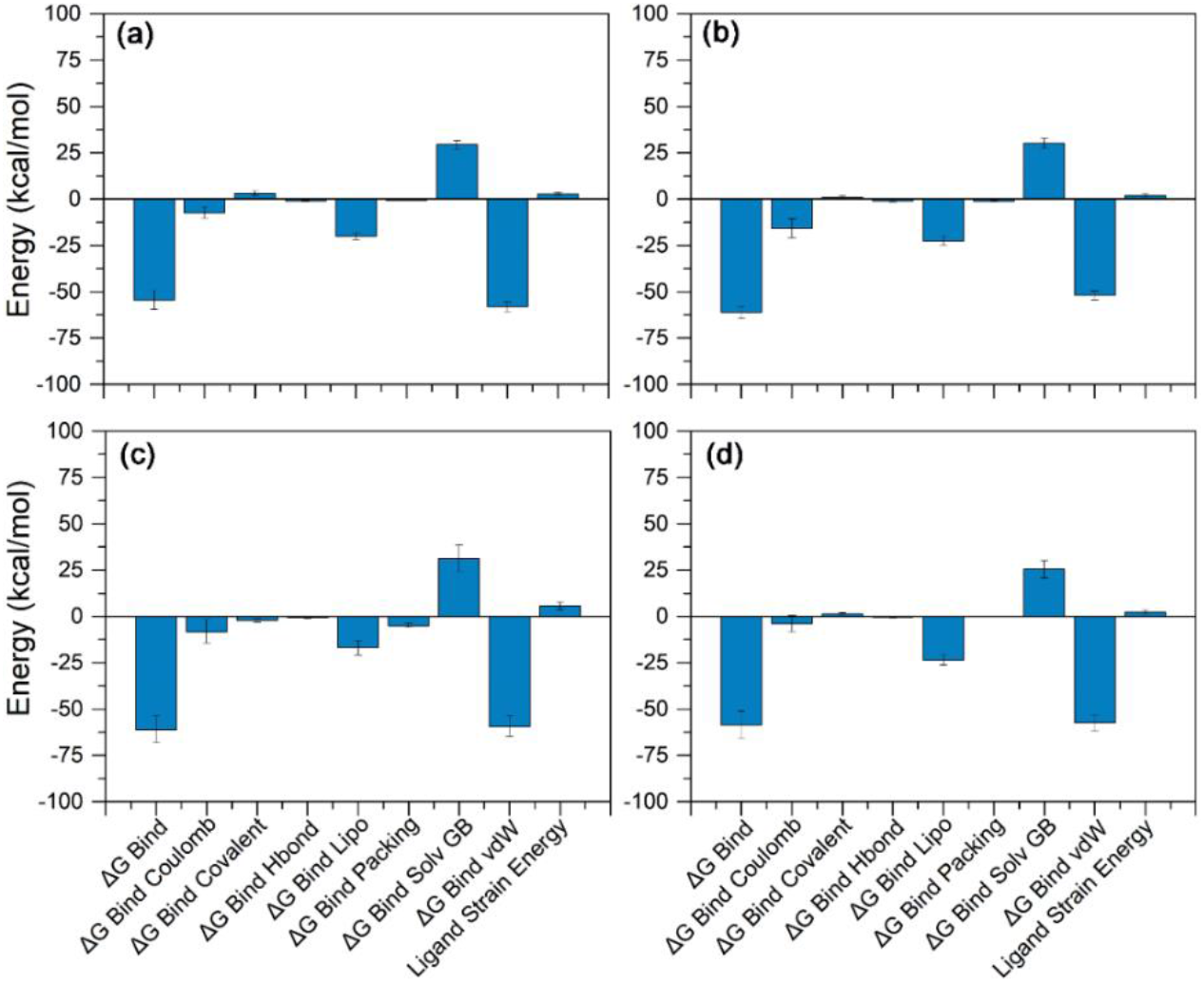
Binding free energy and individual dissociation energy components calculation performed for jRdRp protein docked with bioactive compounds, i.e. (a) Gedunin, (b) Nimbolide, (c) Ohchinin acetate, and (d) Kulactone, on the extracted poses from total 100 ns simulation interval.

## Notes

### Competing Interest Statement

The authors have declared no competing interest.

