## Supplementary material for "Mechanistic insights into the Japanese Encephalitis Virus RNA dependent RNA polymerase protein inhibition by bioflavonoids from *Azadirachta indica*": Table S1, Table S2, Table S3, Table S4, Figure S1, Figure S2, Figure S3, Figure S4, Figure S5, Figure S6, Figure S7

**Results and Discussion**

**S1.1 Structure based virtual screening**

**Table S1.** List of virtual screened bioflavonoids from Azadirachta indica against Japanese Encephalitis Virus RNA

| Compound | Model ID | Energy | nRot |
|---|---|---|---|
| Gedunin | 1 | -11.6 | 0 |
| Nimbolide | 1 | -11 | 0 |
| Ohchinin Acetate | 1 | -11 | 10 |
| Kulactone | 1 | -10.8 | 3 |
| Limocinin | 1 | -10.6 | 6 |
| Nimbinin | 1 | -10.6 | 0 |
| Azadirachtin | 1 | -10.4 | 13 |
| Rutin | 1 | -10.2 | 16 |
| Myricetin | 1 | -10.1 | 0 |
| Azadirachtol | 1 | -10 | 9 |
| Vilasinin | 1 | -10 | 4 |
| Desacetylsalannin | 1 | -9.9 | 8 |
| Nimocinol | 1 | -9.8 | 2 |
| Isomeldenin | 1 | -9.8 | 4 |
| Azadiradione | 1 | -9.8 | 3 |
| Kaempferol-3-O-rutinoside | 1 | -9.7 | 15 |
| Salannin | 1 | -9.6 | 9 |
| Desacetylnimbin | 1 | -9.5 | 7 |
| SALANNOLIDE | 1 | -9.5 | 10 |
| Khivorin | 1 | -9.5 | 7 |
| Kaempferol-3-O-Beta-D-glucoside | 1 | -9.5 | 10 |
| Salannol | 1 | -9.3 | 9 |
| Hyperoside | 1 | -9.2 | 12 |
| Nimbandiol | 1 | -9.1 | 5 |
| nimbidiol | 1 | -9.1 | 5 |
| Nimbin | 1 | -9.1 | 8 |
| Isonimbinolide | 1 | -9.1 | 9 |
| Nimbione | 1 | -9 | 1 |
| Zafaral | 1 | -8.8 | 6 |
| BETA-SITOSTEROL | 1 | -8.8 | 7 |
| Margocinin | 1 | -8.8 | 4 |
| Nimosone | 1 | -8.7 | 2 |
| OhchinolideB | 1 | -8.7 | 8 |
| Quercetin | 1 | -8.6 | 6 |
| Nimbinone | 1 | -8.6 | 1 |
| Kaempferol | 1 | -8.4 | 5 |
| Nimbiol | 1 | -8.4 | 1 |
| Sugiol | 1 | -8.2 | 2 |
| Scopoletin | 1 | -6.7 | 2 |
| Behenic | 1 | -5.2 | 21 |

dependent RNA polymerase (RdRp) protein.

**S1.2. ADME profiling**

**Table S2**: ADMET profiling for the selected bioactive compound from *Azadirachta indica* as inhibitor against jRdRp protein.

| Properties | Gedunin | Nimbolide | Ohchinin acetate | Kulactone |
|---|---|---|---|---|
| iLOGP | 3.22 | 3.51 | 4.42 | 4.54 |
| XLOGP3 | 4.22 | 2.17 | 4.82 | 6.89 |
| WLOGP | 4.24 | 3.74 | 5.93 | 7.06 |
| MLOGP | 2.56 | 2.28 | 3.35 | 5.63 |
| Silicos-IT Log P | 4.44 | 3.83 | 5.79 | 6.72 |
| Consensus Log P | 3.74 | 3.11 | 4.86 | 6.17 |
| ESOL Log S | -5.4 | -3.94 | -6.39 | -6.79 |
| ESOL Solubility (mg/ml) | 1.93E-03 | 5.30E-02 | 2.64E-04 | 7.35E-05 |
| ESOL Solubility (mol/l) | 4.00E-06 | 1.14E-04 | 4.10E-07 | 1.62E-07 |
| ESOL Class | Moderately soluble | Soluble | Poorly soluble | Poorly soluble |
| Ali Log S | -5.93 | -3.74 | -6.87 | -7.61 |
| Ali Solubility (mg/ml) | 5.64E-04 | 8.57E-02 | 8.63E-05 | 1.11E-05 |
| Ali Solubility (mol/l) | 1.17E-06 | 1.84E-04 | 1.34E-07 | 2.44E-08 |
| Ali Class | Moderately soluble | Soluble | Poorly soluble | Poorly soluble |
| Silicos-IT LogSw | -5.75 | -5.27 | -7.65 | -6.7 |
| Silicos-IT Solubility (mg/ml) | 8.50E-04 | 2.49E-03 | 1.44E-05 | 9.03E-05 |
| Silicos-IT Solubility (mol/l) | 1.76E-06 | 5.35E-06 | 2.23E-08 | 2.00E-07 |
| Silicos-IT class | Moderately soluble | Moderately soluble | Poorly soluble | Poorly soluble |
| GI absorption | High | High | Low | Low |
| BBB permeant | No | No | No | No |
| Pgp substrate | Yes | Yes | Yes | No |
| CYP1A2 inhibitor | No | No | No | No |
| CYP2C19 inhibitor | No | No | No | No |
| CYP2C9 inhibitor | No | No | Yes | Yes |
| CYP2D6 inhibitor | No | No | No | No |
| CYP3A4 inhibitor | No | No | No | No |
| log Kp (cm/s) | -6.25 | -7.61 | -6.81 | -4.17 |
| Lipinski #violations | 0 | 0 | 1 | 1 |
| Ghose #violations | 1 | 0 | 4 | 3 |
| Veber #violations | 0 | 0 | 0 | 0 |
| Egan #violations | 0 | 0 | 1 | 1 |
| Muegge #violations | 0 | 0 | 1 | 1 |
| Bioavailability Score | 0.55 | 0.55 | 0.55 | 0.55 |
| PAINS #alerts | 0 | 0 | 0 | 0 |
| Brenk #alerts | 2 | 2 | 3 | 1 |
| Leadlikeness #violations | 2 | 1 | 3 | 2 |
| Synthetic Accessibility | 6.48 | 6.07 | 7.2 | 5.86 |

### S1.3. Re-docking and intermolecular interaction analysis

**Table S3:** List of selected bioflavonoids as inhibitors of viral RdRp protein and molecular interaction profiling in the respective docked complexes.

| S. no. | Compounds | Re-docking Score (kcal/mol) | H-bond | π-π stacking/ *Salt bridge | Hydrophobic | Polar | Negative | Positive | Glycine |
|---|---|---|---|---|---|---|---|---|---|
| 1. | Gedunin | - 10.4 | Ser$^{604}$, Ile$^{802}$ | -- | Leu$^{411}$, Ala$^{413}$, Val$^{414}$, Ala$^{475}$, Ile$^{476}$, Trp$^{477}$, Tyr$^{610}$, Trp$^{800}$, Ile$^{802}$ | Ser$^{604}$, Thr$^{609}$, Asn$^{613}$, Ser$^{801}$ | Asp$^{541}$, Asp$^{668}$ | Arg$^{460}$, Arg$^{474}$ | Gly$^{412}$, Gly$^{605}$ |
| 2. | Nimbolide | -10.9 | Ser$^{604}$, Ser$^{801}$, Ile$^{802}$ | -- | Leu$^{411}$, Ala$^{413}$, Val$^{607}$, Tyr$^{610}$, Cys$^{714}$, Trp$^{800}$, Ile$^{802}$ | Ser$^{604}$, Gln$^{606}$, Thr$^{609}$, Asn$^{613}$, Ser$^{666}$, Ser$^{801}$, His$^{803}$ | Asp$^{668}$, Asp$^{669}$ | Lys$^{404}$, Arg$^{474}$, Lys$^{471}$ | Gly$^{412}$, Gly$^{667}$ |
| 3. | Ohchinin acetate | -11.0 | Trp$^{477}$, Ser$^{604}$, Ser$^{801}$, Ile$^{802}$ | -- | Ala$^{410}$, Leu$^{411}$, Ala$^{413}$, Val$^{414}$, Ala$^{475}$, Ile$^{476}$, Trp$^{477}$, Val$^{607}$, Tyr$^{610}$, Cys$^{714}$, Trp$^{800}$, Ile$^{802}$ | Asn$^{495}$, Ser$^{604}$, Thr$^{609}$, Asn$^{613}$, Ser$^{801}$, His$^{803}$ | Asp$^{668}$, Asp$^{669}$ | Lys$^{404}$, Arg$^{460}$, Arg$^{474}$, Lys$^{471}$ | Gly$^{412}$, Gly$^{603}$, Gly$^{605}$, Gly$^{667}$ |
| 4. | Kulactone | -10.4 | Asn$^{613}$ | -- | Val$^{414}$, Phe$^{415}$, Ala$^{475}$, Ile$^{476}$, Phe$^{478}$, Tyr610, Trp$^{800}$ | Thr$^{346}$, Ser$^{604}$, Thr$^{609}$, Asn$^{613}$, Ser$^{666}$ | Asp$^{541}$, Asp$^{668}$ | Arg$^{460}$, Arg$^{474}$ | Gly$^{412}$, Gly$^{605}$, Gly$^{667}$ |
| 5. | Guanosine-5'-triphosphate | -9.0 | Lyn$^{463}$, Arg$^{474}$, Asp$^{668}$, Asp$^{669}$(2), Trp$^{800}$, Ile$^{802}$ | *Arg$^{474}$(3), ***Arg$^{460}$ (2)**, *Arg$^{742}$ | Met$^{345}$, Tyr$^{610}$, Cys$^{714}$, Trp$^{800}$, Ile$^{802}$ | Ser$^{604}$, Ser$^{666}$, Ser$^{715}$, Ser$^{801}$, His$^{803}$ | Asp$^{668}$, Asp$^{669}$ | Arg$^{460}$, Lys$^{471}$, Arg$^{474}$, Arg$^{742}$ | Gly$^{412}$, Gly$^{667}$ |

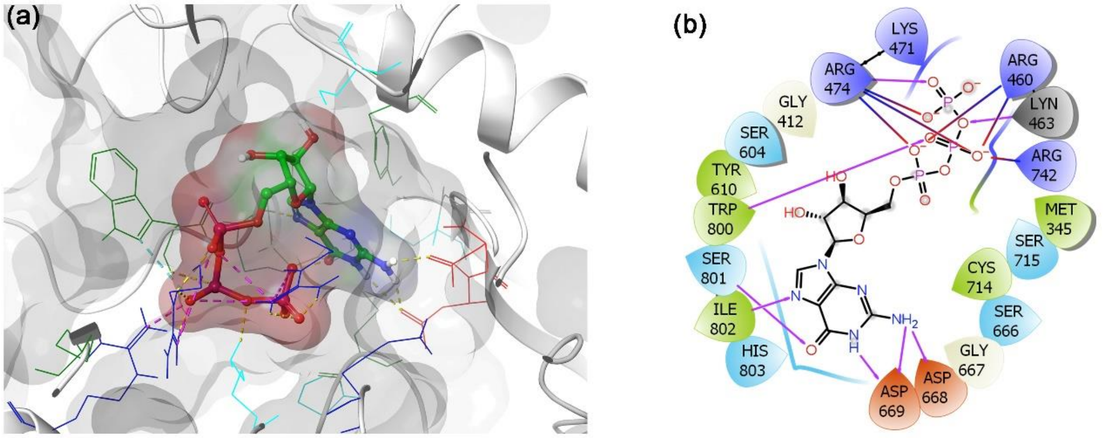


**Figure S1**. 3D and 2D docked poses of the jRdRp-GTP docked complex collected at 4 Å space around the ligand within in the active site of jRdRP protein. In 3D structures, protein surface and ligand surface were rendered based on the alpha-carbon and atomic charge, respectively. While in 2D maps, hydrogen bond formation (pink arrows), Salt bridge (red-violet lines) hydrophobic (green), polar (blue), red (negative), violet (positive), glycine (grey) interactions are logged for docked complexes of jRdRp with selected bioactive compounds.

**S1.4. Classical molecular dynamics simulation analysis**

**S1.4.1.** ***RMSD and RMSF analysis***

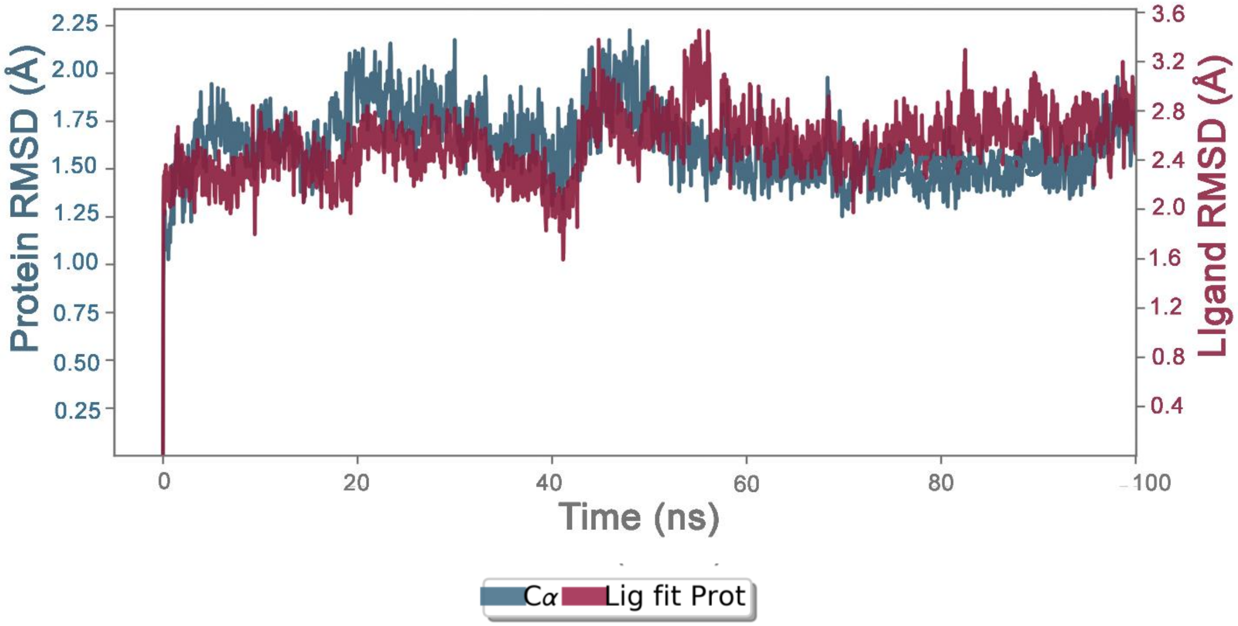


**Figure S2:** RMSD plot for the backbone atoms of RdRp in complex with the reference inhibitor, i.e. GTP.

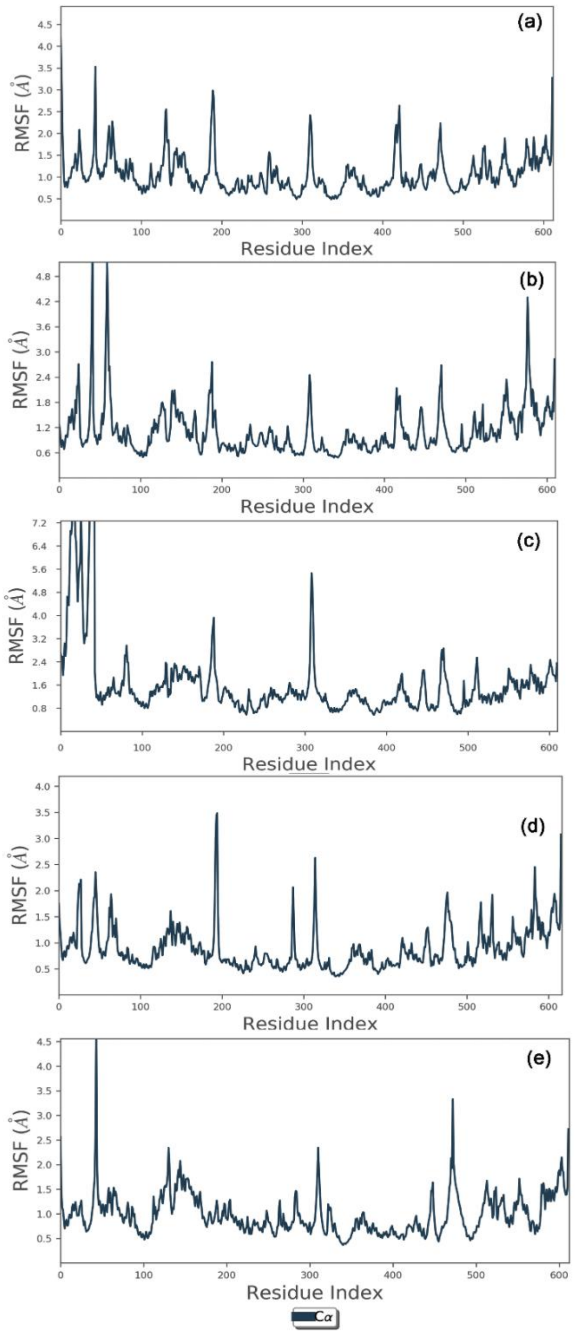


**Figure S3.** RMSF plot generated for the jRdRp docked with selected bioactive compounds, i.e., i.e., (a) Gedunin, (b) Nimbolide, (c) Ohchinin acetate, and (d) Kulactone, and reference ligand, viz, (e) GTP, during 100 ns molecular dynamics simulation interval. Herein, residue number 0-612 are actually residue number from 274-889.

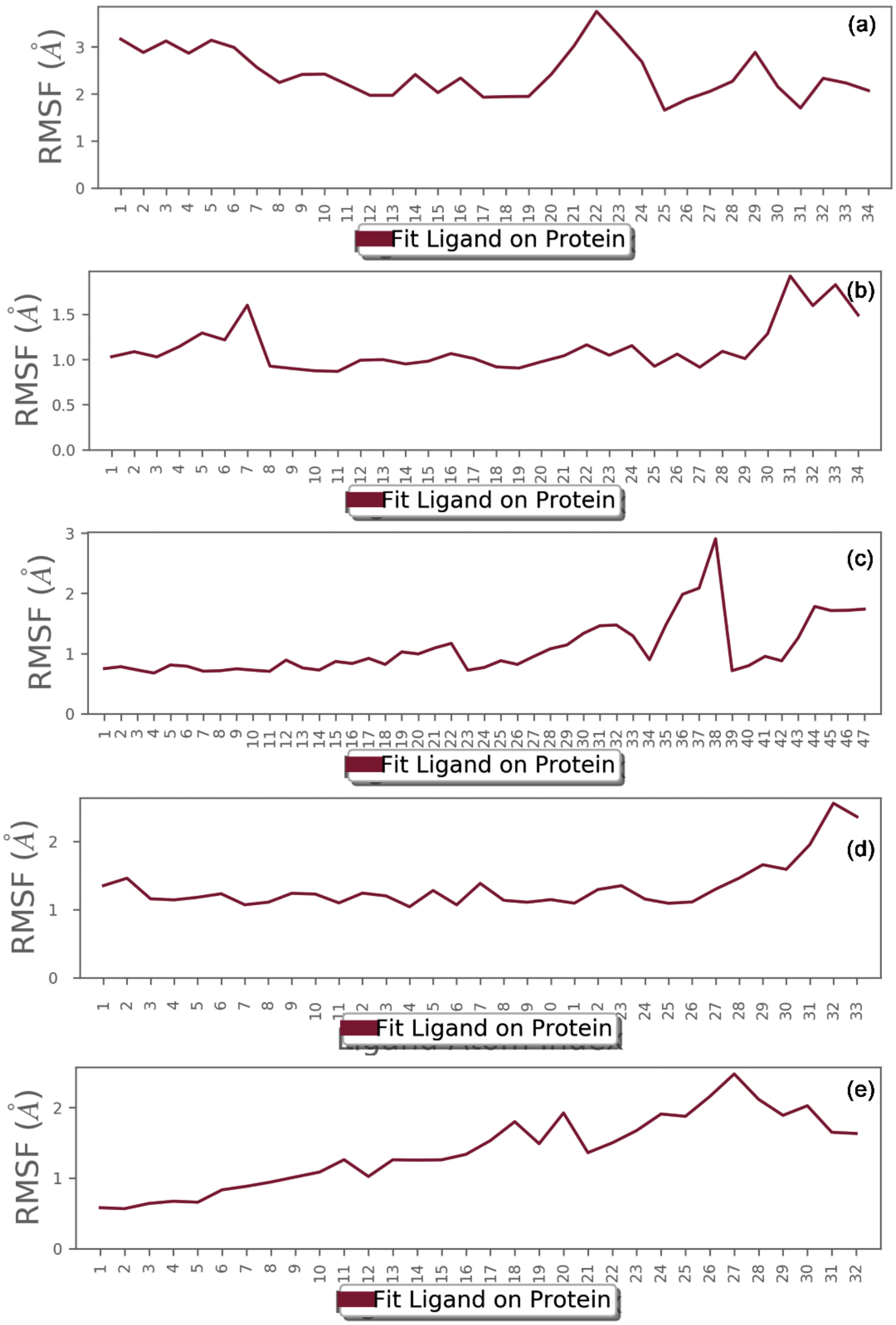


**Figure S4.** RMSF plot generated for the docked bioactive compounds, i.e., (a) Gedunin, (b) Nimbolide, (c) Ohchinin acetate, and (d) Kulactone, and reference ligand, viz, (e) GTP, fit in the jRdRp protein during 100 ns molecular dynamics simulation interval.

**S1.4.2.** ***Protein-ligand interaction profiling***

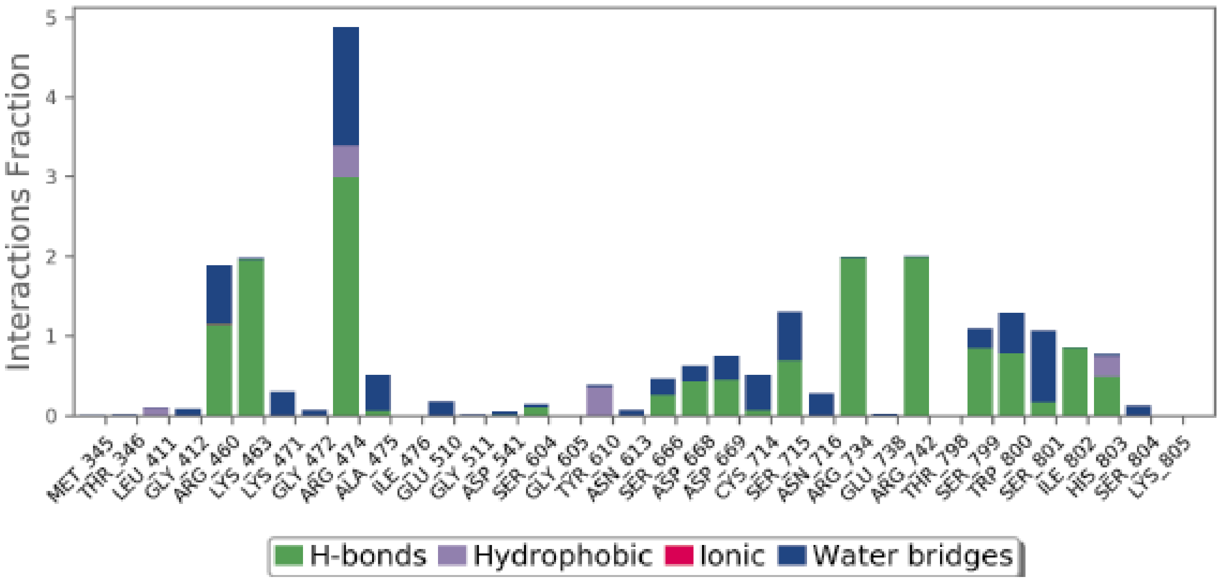


**Figure S5.** Protein-ligand interactions mapping for jRdRp docked with reference compound, i.e. GTP, extracted from 100 ns MD simulations. Herein, values of interaction fractions > 1.0 are feasible as some residues established several interactions of the similar subtype.

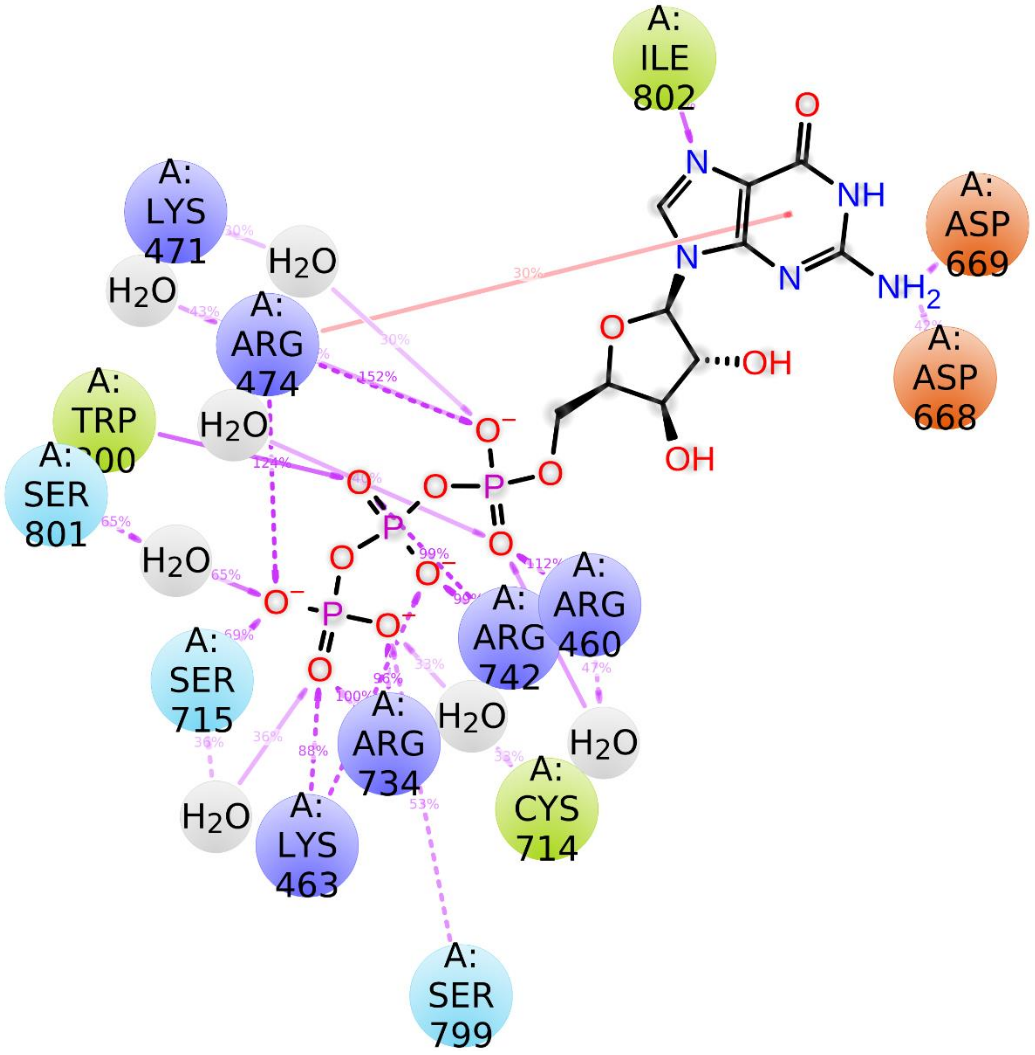


**Figure S6.** Schematic representation for interaction profile of jRdRp-GTP complex extracted at 30% of total 100 ns simulation interval.

### S1.5. Binding free energy analysis

**Table S4** : Binding free energy and individual dissociation energy components calculated for selected docked complexes of jRdRp protein with bioactive compounds, i.e. (a) Gedunin, (b) Nembolide, (c) Ohchinin acetate, and (d) Kulactone.

| MM/GBSA Components | Energy (kcal/mol) | | | | |
|---|---|---|---|---|---|
| | jRdRp-Gedunin | jRdRp-Nembolide | jRdRp-Ohchinin acetate | jRdRp-Kulactone | jRdRp-GTP |
| ΔG Bind | -54.54±5.09 | -61.13±3.26 | -61.09±7.08 | -58.65±7.41 | -78.9±9.05 |
| $\Delta G_{Bind\ Coulomb}$ | -7.27±3.05 | -15.73±5.14 | -8.26±6.35 | -3.96±4.32 | -217.04±21.49 |
| $\Delta G_{Bind\ Covalent}$ | 3.03±1.23 | 1.15±0.57 | -2.1±1.13 | 1.24±0.68 | 6.79±2.61 |
| $\Delta G_{Bind\ Hbond}$ | -0.95±0.35 | -0.96±0.54 | -0.54±0.61 | -0.37±0.41 | -17.83±2.05 |
| $\Delta G_{Bind\ Lipo}$ | -20.03±1.91 | -22.57±2.27 | -16.98±3.91 | -23.5±2.85 | -6.62±0.76 |
| $\Delta G_{Bind\ Packing}$ | -0.6±0.22 | -1.14±0.32 | -5.03±1.17 | 0±0 | -3.12±0.81 |
| $\Delta G_{Bind\ Solv\ GB}$ | 29.36±2.21 | 30.09±2.8 | 31.24±7.04 | 25.45±4.57 | 203.73±14.99 |
| $\Delta G_{Bind\ vdW}$ | -58.09±2.68 | -51.95±2.27 | -59.42±5.37 | -57.48±4.31 | -44.83±4.41 |
| Lig Strain Energy | 2.86±0.73 | 1.97±0.8 | 5.56±1.98 | 2.19±1.1 | 6.98±4.03 |

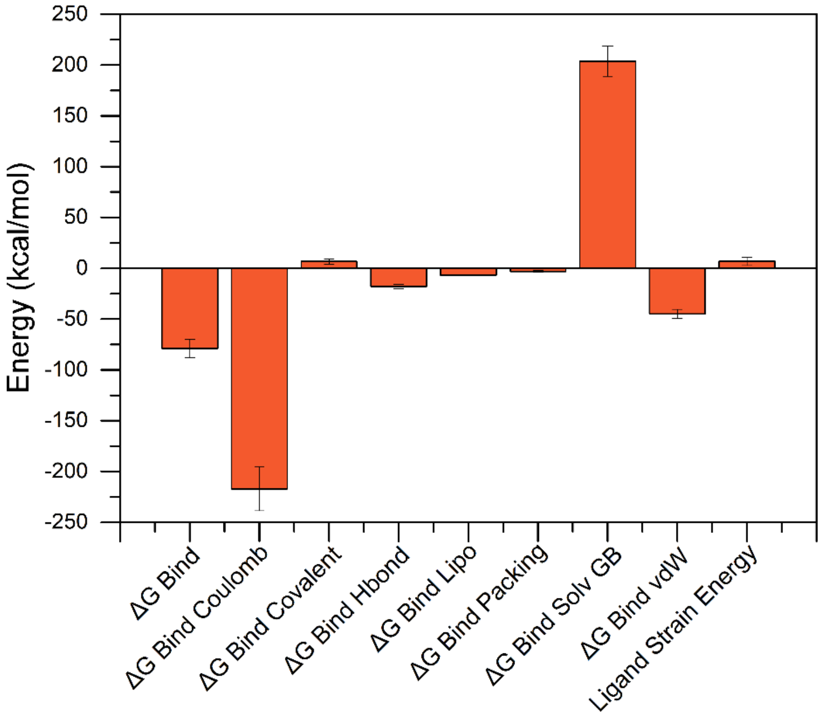


**Figure S7**. Binding free energy and individual dissociation energy components calculation performed for the reference jRdRp-GTP complex.
